# REViewer: Haplotype-resolved visualization of read alignments in and around tandem repeats

**DOI:** 10.1101/2021.10.20.465046

**Authors:** Egor Dolzhenko, Ben Weisburd, Kristina Ibanez Garikano, Indhu Shree Rajan Babu, Mark F Bennett, Kimberley Billingsley, Ashley Carroll, Matt C. Danzi, Viraj Deshpande, Jinhui Ding, Sarah Fazal, Andreas Halman, Bharati Jadhav, Yunjiang Qiu, Phillip Richmond, Konrad Scheffler, Joke J.F.A van Vugt, Ramona R.A.J. Zwamborn, Genomics England Research Consortium, Samuel S. Chong, Jan M. Friedman, Arianna Tucci, Heidi L. Rehm, Michael A Eberle

## Abstract

**Background:** Expansions of short tandem repeats are the cause of many neurogenetic disorders including familial amyotrophic lateral sclerosis, Huntington disease, and many others. Multiple methods have been recently developed that can identify repeat expansions in whole genome or exome sequencing data. Despite the widely-recognized need for visual assessment of variant calls in clinical settings, current computational tools lack the ability to produce such visualizations for repeat expansions. Expanded repeats are difficult to visualize because they correspond to large insertions relative to the reference genome and involve many misaligning and ambiguously aligning reads.

**Results:** We implemented REViewer, a computational method for visualization of sequencing data in genomic regions containing long repeat expansions. To generate a read pileup, REViewer reconstructs local haplotype sequences and distributes reads to these haplotypes in a way that is most consistent with the fragment lengths and evenness of read coverage. To create appropriate training materials for onboarding new users, we performed a concordance study involving 12 scientists involved in STR research. We used the results of this study to create a user guide that describes the basic principles of using REViewer as well as a guide to the typical features of read pileups that correspond to low confidence repeat genotype calls. Additionally, we demonstrated that REViewer can be used to annotate clinically-relevant repeat interruptions by comparing visual assessment results of 44 *FMR1* repeat alleles with the results of triplet repeat primed PCR. For 38 of these alleles, the results of visual assessment were consistent with triplet repeat primed PCR.

**Conclusions:** Read pileup plots generated by REViewer offer an intuitive way to visualize sequencing data in regions containing long repeat expansions. Laboratories can use REViewer to assess the quality of repeat genotype calls as well as to visually detect interruptions or other imperfections in the repeat sequence and the surrounding flanking regions.

## Background

Visual inspection of sequencing data supporting a given genetic variant is an important part of clinical bioinformatics pipelines. Effective visualizations enable scientists to quickly assess the quality of sequencing data supporting a genotype call. Factors that impact genotyping accuracy such as local depth, evenness of coverage, presence of any additional variation, and other locus-specific features are difficult to piece together from genome-wide quality metrics and various per-variant scores typically reported by variant calling methods. Recent guidelines from the Association for Medical Pathology and the College of American Pathologists strongly recommend review of such visualizations during routine sign out of variant calls (Roy et al. 2018).

The Integrative Genomics Viewer (Robinson et al. 2011), JBrowse (Buels et al. 2016), and other general-purpose tools for visualization of sequencing data work well for single nucleotide variants, short indels, and copy number variants. Additionally, specialized methods have been developed for visualizing reads associated with variants that involve more complex indel patterns and distal breakpoints (Gymrek 2014; Nattestad et al. 2021; Spies et al. 2015; Belyeu et al. 2021). However, there is a lack of methods for visualizing sequencing data in regions harboring long repetitive sequences such as long stretches of short tandem repeats (STRs).

Analysis and visualization of regions containing long STRs using short read sequencing data pose a number of unique challenges. For instance, it is difficult to correctly align reads originating within the sequence of a long STR because the number of possible alignment positions increases linearly with the length of the STR allele. Regions containing multiple adjacent STRs—including the regions linked with Huntington disease, Friedreich ataxia, and Spinocerebellar ataxia 8—are especially prone to alignment artifacts because adjacent repeats may have a high sequence similarity and because the sizes of these repeats in a given individual often differ from those in the reference genome.

Here we present the Repeat Expansion Viewer (REViewer), a novel method for visualizing short read sequencing data in genomic regions containing one or multiple STRs. REViewer has been designed to work with the read alignments produced by ExpansionHunter (Dolzhenko et al. 2017, 2019), though it will work with any repeat genotyping software that produces output in the appropriate format. We also describe FlipBook, a companion image viewer that is designed for manual curation of large collections of images generated by REViewer.

## Implementation

### Overview

REViewer is designed to work with the BAM (Li et al. 2009) and VCF files (Danecek et al. 2011) generated by ExpansionHunter (Dolzhenko et al. 2017, 2019), a commonly used method for repeat genotyping. The VCF file is used to obtain repeat genotypes while the BAM file contains reads realigned to a sequence graph representing the entire repeat region (**Figure 1**, panels 1-4). Additionally, we created a wrapper script that accepts regular BAM files containing alignments of reads to a linear genome and a tab-separated file containing reference coordinates of the target STRs, repeat units, and repeat genotypes making it possible to use REViewer with other software (**supplementary information**).

**Figure 1:**
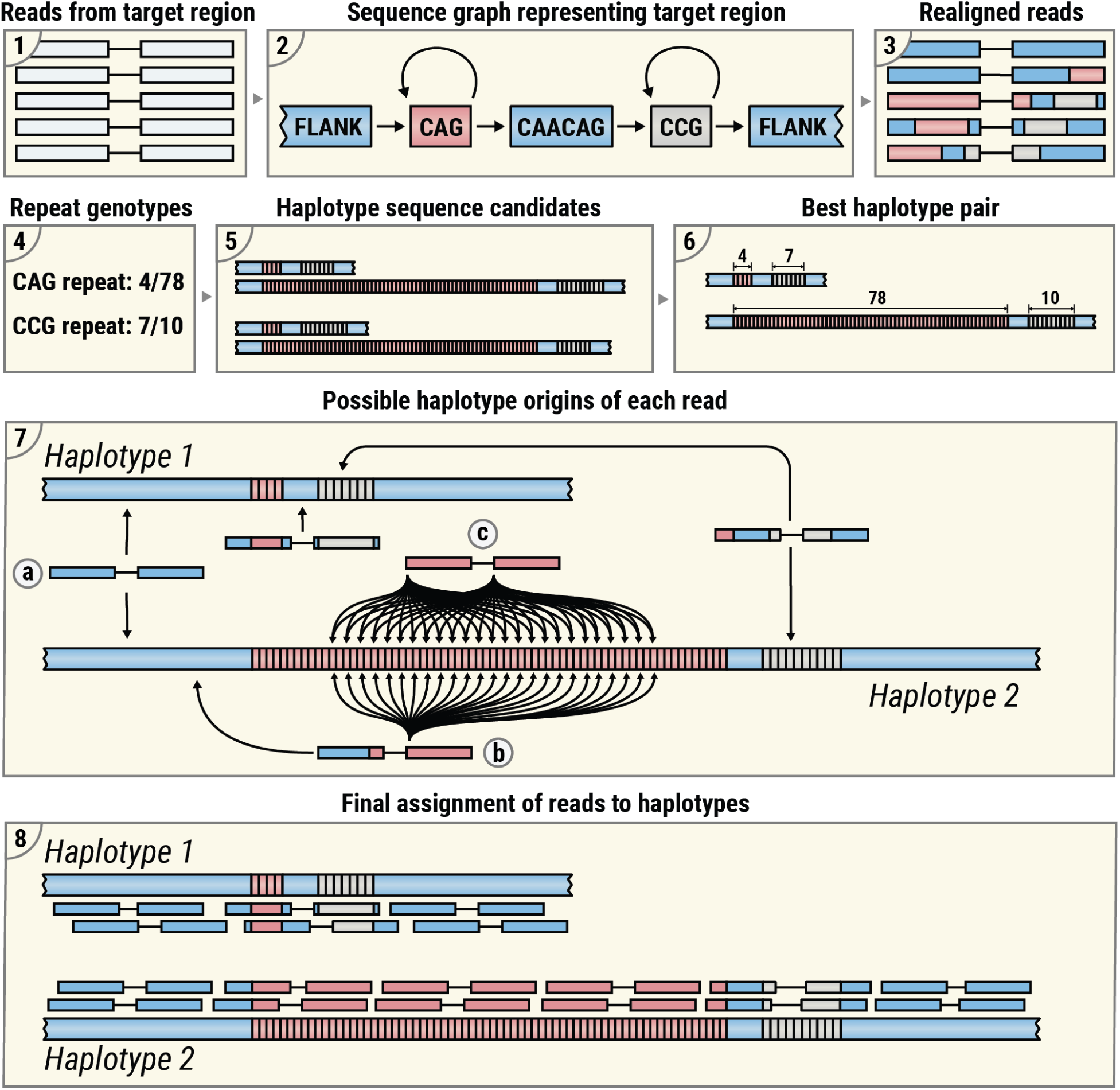
An overview of the pileup generation algorithm: (1-3) reads originating in the region containing target STRs are realigned using the sequence graph aligner within ExpansionHunter software; (4,5) putative pairs of haplotype sequences are generated from repeat genotypes; (6) a haplotype pair that has the highest consistency with read alignments is selected; (7) possible alignments of each read to each haplotype sequence are generated from the original sequence-graph alignments; (8) pairs of read alignments that correspond to the most consistent fragment length are selected for each read pair and then one of these is randomly selected for visualization.

### Read pileup generation

Read pileups are generated using genotypes of all STRs present at the target region and reads aligned to a sequence graph representing the region (**Figure 1**, panel 1-4; (Dolzhenko et al. 2019)). For repeats on diploid chromosomes, REViewer constructs all possible pairs of haplotype sequences from the STR genotypes. For example, if a region contains two STRs then there are four possible haplotypes that can be formed and two possible haplotype pairings (**Figure 1**, panel 5). The reads are next aligned to all haplotype pairs by transforming the graph alignments from the BAM file generated by ExpansionHunter into linear alignments. The haplotype pair that yields the highest cumulative read alignment score is selected for visualization (**Figure 1**, panel 6). Loci with a single STR or on haploid chromosomes have unambiguous haplotypes and so the haplotype sequence selection steps are skipped. Next, for each read pair, REViewer finds the top-scoring alignments to any haplotype sequence (**Figure 1**, panel 7). A read pair originating completely within a sequence surrounding the repeats and shared by all haplotypes has exactly one alignment position on each haplotype (**Figure 1**, panel 7a). When one mate originates fully within the repeat, the number of positions for the read pair increases linearly with the repeat length (**Figure 1**, panel 7b). In contrast, when both mates originate inside the repeat, the number of positions increases quadratically (**Figure 1**, panel 7c). For read pairs where one or both mates have multiple alignments, REViewer selects pairs of alignments that correspond to fragment length closest to the mean fragment length calculated for read pairs mapping to the flanking regions surrounding the repeats. Finally, REViewer generates read pileup by selecting one pair of alignments at random for each read pair (**Figure 1**, panel 8).

This algorithm is based on the idea that if a given locus is sequenced well and each constituent repeat is genotyped correctly, then it is possible to distribute the reads to achieve an even coverage of each haplotype. Importantly, assignment of some reads to the correct haplotype of origin will be ambiguous, especially in cases when the repeats are homozygous, and the resulting haplotypes are identical.

Pileups corresponding to correctly genotyped repeats are characterized by a relatively even read coverage of both alleles (**Figure 2**, panels 1-3). At the typical whole-genome sequencing depths (30-60x), each position of a haplotype sequence is expected to be covered by many reads (15-30), though the coverage may dip in certain regions due to technical factors like GC bias. For repeats much shorter than the read length, this implies the presence of multiple spanning reads (**Figure 2**, both alleles on panel 1 and short allele on panel 2). Repeats much larger than the read length are expected to contain multiple in-repeat reads (**Figure 2**, long allele on panel 1 and both alleles on panel 2). An incorrectly called expanded allele might have low sequencing depth inside the repeat compared to the depth of the region surrounding the repeat (**Figure 2**, long allele on panel 4). Additionally, the presence of multiple indels in the alignments of in-repeat reads indicates that the reads may not be correctly aligned (possibly due to sequencing errors) and that the size of the repeat may be overestimated (**Figure 2**, panel 5).

**Figure 2:**
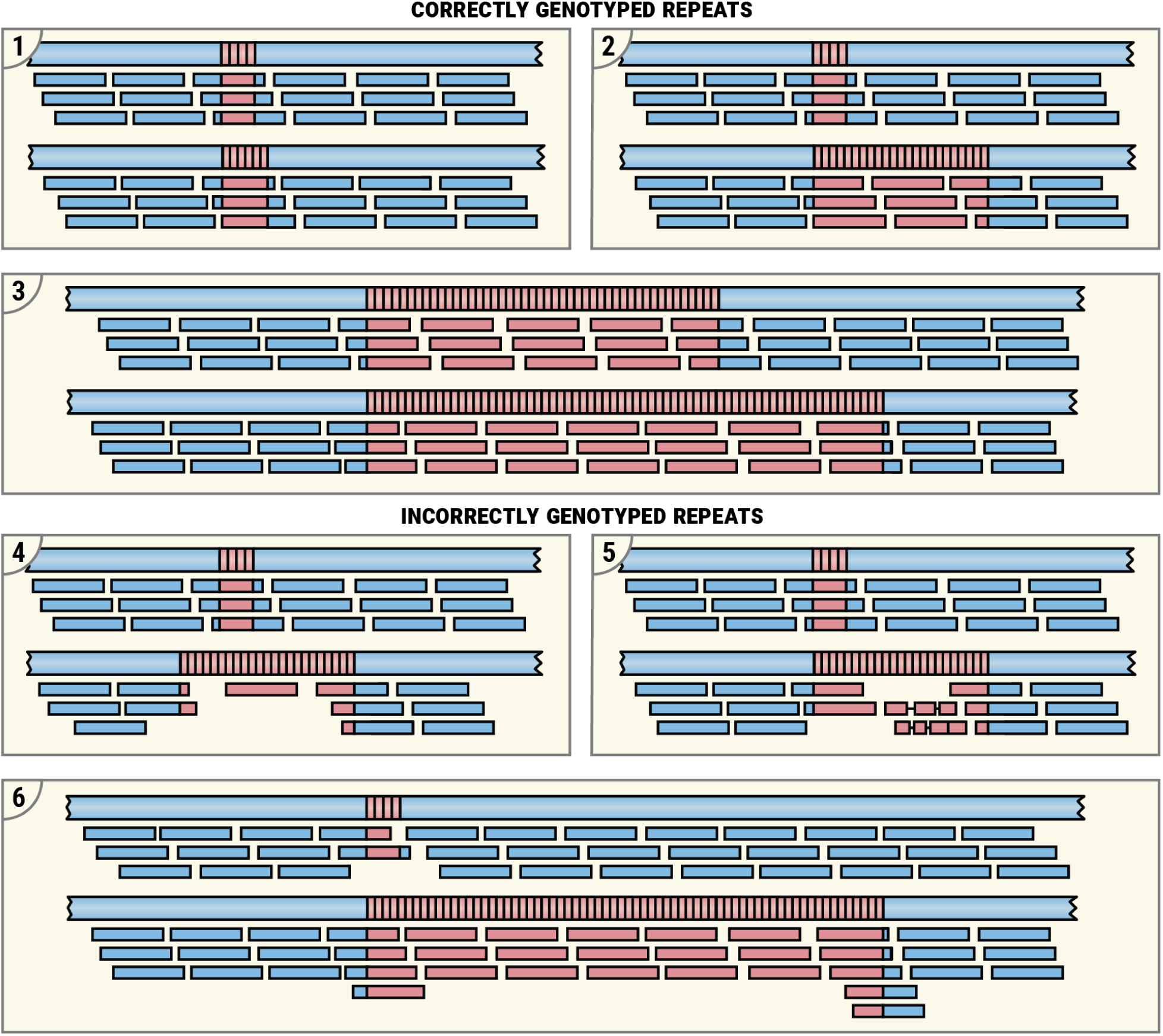
Examples of read pileups. Pileups corresponding to correctly genotyped repeats: (1) both repeat alleles are short; (2) one allele is expanded; (3) both alleles are expanded. Pileups corresponding to incorrectly genotyped repeats: (4) expanded allele is supported by just one read suggesting that its size is overestimated; (5) expanded allele is supported by poorly aligning reads (each containing multiple indels) suggesting that the reads are incorrectly mapped and that size of the repeat is overestimated; (6) the short allele is supported by just one spanning read suggesting that this allele is not real and that both alleles are expanded.

Finally, a short allele supported by one or very few spanning reads may not be real. For instance, the short allele depicted on panel 6 of **Figure 2** is supported by just one spanning and one flanking read, which is less than expected based on the coverage of the surrounding region. There is also a slight excess of the flanking reads on the long allele of this repeat. Taken together, these observations suggest that (a) the single spanning read may be a result of an incorrect alignment and (b) the correct genotype is likely to be a double expansion. Some real examples corresponding to the scenarios depicted in **Figure 2** are included in online documentation (*Examples*.*md at Master · Illumina/REViewer* n.d.).

### FlipBook image viewer

In many situations, researchers may wish to look at STR genotypes for a variety of known repeat loci across many samples. To simplify the painstaking manual task of reviewing many REViewer pileups and recording the results of manual review, we developed FlipBook—a photo-album-like application that lets a user quickly assess pileups on their local hard drive and record notes about each one. Additional features of this software include 1) displaying custom information above the images—such as affected status and STR locus information, 2) customizing the questions a user can answer about each image, and 3) displaying more than one image at a time—such as when evaluating data from multiple family members.

## Results

### A concordance study

To solicit feedback on REViewer and FlipBook and create training materials for new REViewer users, we performed a concordance study involving 12 scientists (analysts). We used a collection of whole-genome sequencing (WGS) samples described in a recent study of subjects with suspected neurological disorders (Ibanez et al. 2020) and additional samples from the 100,000 Genomes Project that were validated by PCR (see **supplementary information**). The *HTT, TBP, AR, ATXN3, ATN1, ATXN2, ATXN7, ATXN1, CACNA1A, DMPK, PPP2R2B, FXN, FMR1*, and *C9orf72* STR loci were genotyped in these samples with ExpansionHunter (EH) and also tested with PCR. To emulate a practical assessment strategy, only the STRs for which the size confidence interval reported by EH overlapped or exceeded an intermediate or full expansion threshold were selected for review. This totaled 133 STR genotypes (one genotype per sample) across all 14 STR loci. REViewer read pileups corresponding to these 133 genotypes (**Table S1**) were evaluated by the analysts using FlipBook software. The analysts categorized the genotyped STRs into normal, intermediate expansion, full expansion, and biallelic expansion categories. The verdicts were recorded by FlipBook for subsequent analysis.

To measure consistency of analysts’ responses, we calculated the number of discordant verdicts for each genotyped STR. A verdict was defined as discordant if it differed from the most common consensus verdict. The majority of verdicts were highly consistent—three or more analysts disagreed with the consensus verdict for only 9 out of 133 genotyped STRs (**Figure 3**, panel 1). The mean number of STRs with discordant verdicts was below one for all STR loci (**Figure 3**, panel 2). *FMR1* repeats had the largest number of discordant verdicts (0.94 on average) which is consistent with earlier observations that the *FMR1* locus is harder to size accurately as the repeat becomes long (Dolzhenko et al. 2017). Disagreements in verdicts arose for STRs where the size estimate was close to the pathogenic threshold (**Figure 3**, panel 3).

**Figure 3:**
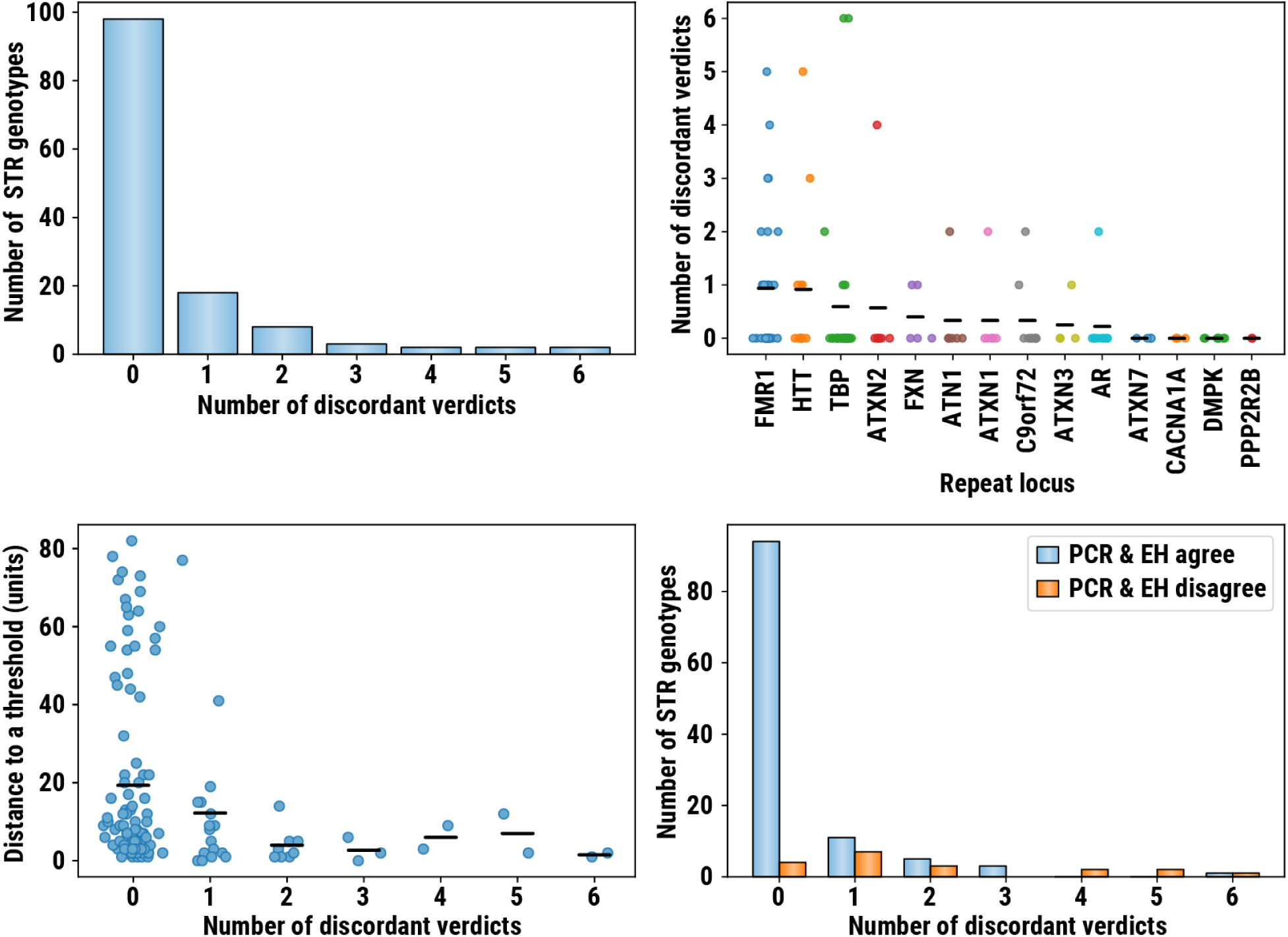
(1) counts of STR genotypes where the specified number of analysts disagreed with the consensus verdict (discordant verdict); (2) distribution of discordant verdict counts stratified by STR locus; (3) distribution of distances between STR sizes estimated by ExpansionHunter (EH) and PCR; (4) the counts of repeat genotypes where NGS and PCR agree and disagree.

Next, we compared the verdicts for repeats where EH and PCR-based calls agreed to those where they disagreed (using binary categorization for *FMR1* and *C9orf72* repeats; see below). When EH and PCR-based calls agreed, most repeats (94 out of 114) had no discordant verdicts but when the EH and PCR-based calls disagreed, only a few (4 out of 19) had no discordant verdicts (**Figure 3**, panel 4). This suggests that, with additional training, the information presented in REViewer/FlipBook visualizations can be used to reduce the false positive rate for many known pathogenic loci. To provide such training, we created online documentation that consists of both a tutorial describing how to review the pileups (**Figure 2** and (*REViewer: A Tool* for Visualizing Alignments of Reads in Regions Containing Tandem Repeats n.d.)) and a repository of pileups corresponding to harder to interpret correct and incorrect calls.

### *FMR1* and *C9orf72* repeat loci

Due to the difficulty of distinguishing between the intermediate and full expansions of *FMR1* (Dolzhenko et al. 2017) *and C9orf72* repeats (full expansions start at 600bp and 360bp respectively), they were categorized into two categories: normal and expanded. This categorization also reflects the fact that, in practice, the ability to distinguish between normal and abnormal sized repeats is more important than being able to accurately classify intermediate versus expanded alleles. Individuals identified with abnormal-sized repeats that may explain their phenotype or place them at risk for disease or passing on an expandable repeat are likely to be sent for orthogonal confirmation testing, regardless of whether the estimated STR size is in the intermediate or pathogenic range.

### Annotating interruptions with REViewer

REViewer visualizations also display deviations from the predicted sequence and this can allow users to identify STR interruptions. To demonstrate this functionality, we assessed the pileups of 29 *FMR1* reference samples (Dolzhenko et al. 2017) with prior TP-PCR data (Rajan-Babu et al. 2015; Chen et al. 2010) on repeat length and number and position of AGG interruptions. The concordance between AGG-interruption maps derived from the REViewer pileups and TP-PCR was evaluated. **Figure 4** shows the read pileups and TP-PCR electropherograms of two representative samples–a normal male (NA06890, panel 1) and an intermediate female (NA20234, panel 2). NA06890 with 30 repeat units has two AGGs evident in the pileups as mismatches at repeat positions 11 and 21. This (CGG)_10_AGG(CGG)_9_AGG(CGG)_9_ structure is consistent with TP-PCR. In NA20234, the pileups show the clear assignment of reads to the correct haplotypes, a 31-repeat normal and a 46-repeat intermediate allele with (CGG)_10_AGG(CGG)_9_AGG(CGG)_10_ and (CGG)_9_AGG(CGG)_9_AGG(CGG)_13_AGG(CGG)_12_ structures, respectively. The TP-PCR analysis had consistent repeat structures, but the superimposing amplicon peaks from the two *FMR1* alleles in some heterozygous female samples with complex repeat structures may make AGG-interruption mapping relatively harder with TP-PCR (Rajan-Babu et al. 2015).

**Figure 4:**
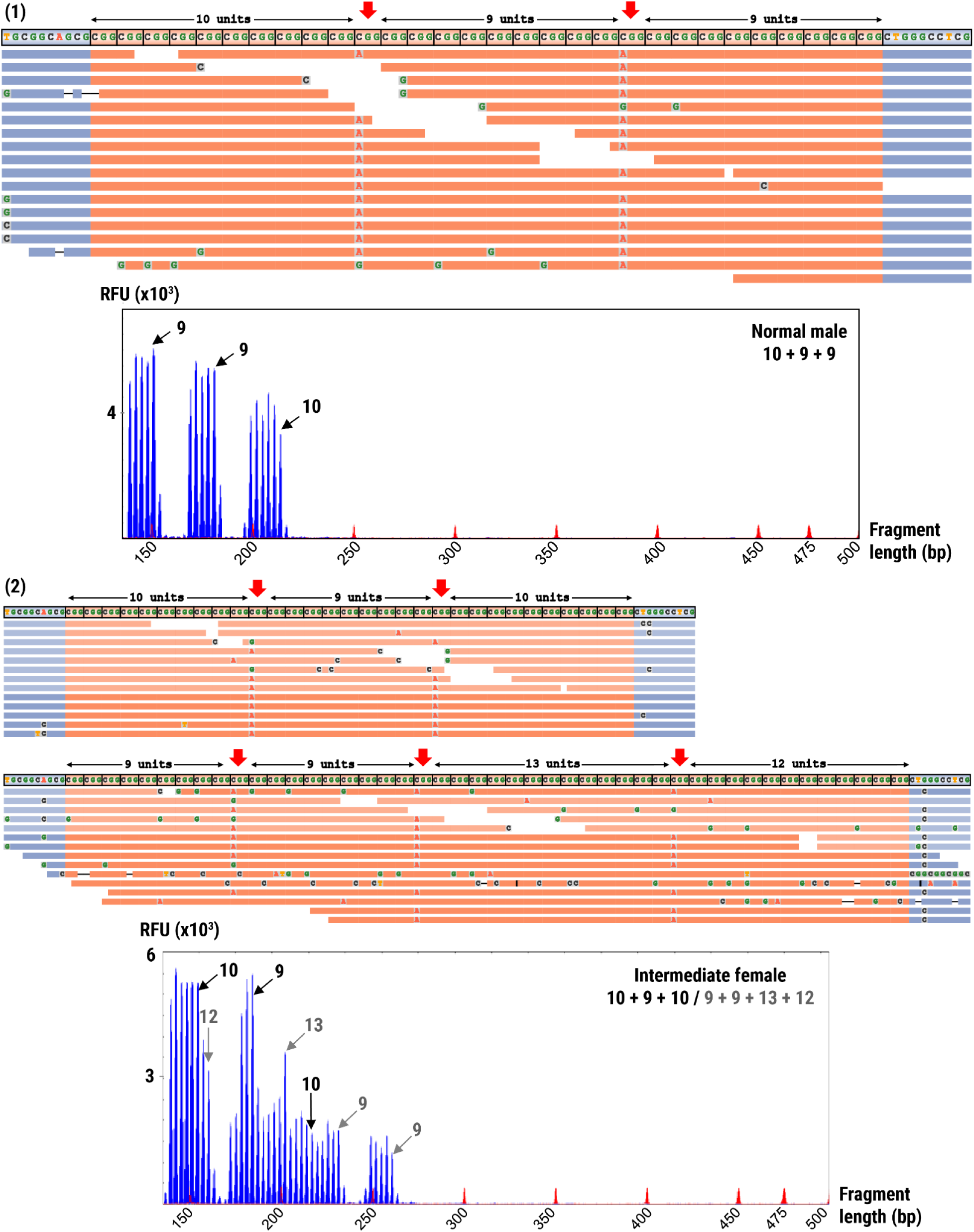
REViewer read pileups and TP-PCR electropherograms of *FMR1* repeats in samples (1) NA06890 and (2) NA20234.

Of the 44 alleles assessed in total (14 males and 15 females), the AGG-interruption maps of 38 alleles derived from the pileups were consistent with that of TP-PCR (**Table S2**). Concordant results (86.36%) were noted for 20/23 normal, 5/5 intermediate, 6/8 premutation, and 7/8 3’-uninterrupted full-mutation alleles.

Among the six discrepant alleles, the normal alleles of NA20243 and NA20240 had an incorrect ExpansionHunter genotype and inadequate spanning/flanking reads in the pileups that hampered the interpretation of AGG interruptions. The normal allele of NA20244 was sized one CGG-repeat less by ExpansionHunter, and the pileup and TP-PCR structures were (CGG)_9_AGG(CGG)_8_AGG(CGG)_21_ and (CGG)_9_AGG(CGG)_9_AGG(CGG)_21_, respectively. We could not resolve the AGG-interruption pattern of the premutation allele in NA20240 due to the ambiguity in the assignment of reads to the two haplotypes as ExpansionHunter genotyped this heterozygous premutation sample (30/80 repeats) as homozygous premutation (95/95 repeats). In NA06907, the premutation haplotype did not have sufficient reads to support the TP-PCR’s (CGG)_10_AGG(CGG)_80_ repeat structure. In NA07537, we could not confidently ascertain the interruption pattern of the full-mutation allele from the pileups because of the ambiguity in read assignment. In general, the TP-PCR data supported the presence of uninterrupted CGG-repeats at the 3’ ends of the full-mutation alleles. Nonetheless, in two full-mutation males (NA06852 and NA06897), the pileup visualization enabled the detection of an AGG interruption at the 5’ end of the full-mutation, which, as expected, was not evident from the TP-PCR analyses that target the 3’ ends. See **Additional File 2** for pileups and TP-PCR profiles of additional *FMR1* intermediate, premutation, and full-mutation samples.

## Discussion

REViewer enables visualization of sequencing data in genomic regions containing one or more tandem repeats by reconstructing local haplotypes containing the repeats of interest and then generating read pileups over these haplotypes. FlipBook, the companion image viewer for REViewer, enables interactive curation of large sets of read pileups and subsequent output of the curation results into a file. We have shown that REViewer and FlipBook can be used for a wide range of purposes including quality assessment of repeat expansion calls produced by bioinformatics pipelines and studies of interruptions and other imperfections in repeat sequences. Additionally, these visualizations are a valuable tool for continued development of new methods for STR analysis.

To create a user guide for REViewer, we performed a concordance study involving 12 scientists involved in STR research. This study highlights a range of pileup features (Figure 2 and (REViewer: A tool for visualizing alig&)) that can help to identify lower confidence calls and potential genotyping errors. This information, together with representative example pileups, was documented in the online user guide. The concordance study also helped to highlight some important limitations of REViewer. Namely, pileups cannot be used to determine if the size of a long repeat expansion is underestimated. This is because pileups of longer repeats missing in-repeat reads can be indistinguishable from pileups corresponding to shorter repeats.

REViewer visualization offers the unique advantage of analysing interruptions at both the 5’- and 3’-ends of the repeat sequences and determining the exact sequences of the interrupting motifs. In the extremely GC-rich *FMR1* repeat locus, which is prone to coverage bias, REViewer achieved an overall 86.36% concordance across normal, intermediate, premutation, and full-mutation genotypes. Interruptions are observed in a number of repeat expansions and their presence or absence may modify the pathogenicity, disease severity or presentation (Matsuura et al. 2006; Kraus-Perrotta and Lagalwar 2016; Cumming et al. 2018). The ability to visualize and assess interruptions is a valuable addition to bioinformatic repeat expansion pipelines. We believe that future improvements to ExpansionHunter genotyping and REViewer’s ability to consider interruptions during the assignment of reads to the haplotypes will enable even better annotations of STR interruptions.

We are planning to continue improving REViewer and FlipBook in response to feedback from the user community. In particular, we are considering extending REViewer to support other variant types.

## Conclusions

Clinical applications of sequencing data continue to rapidly expand. Bioinformatics pipelines for genome analysis continue to increase the types of variants that they profile and incorporate even more difficult regions of the genome. Visualization of sequencing evidence supporting more complex variants requires specialized visualization algorithms and user interfaces. The work here demonstrates that variant-specific visualizations that augment general purpose visualization tools are a pragmatic strategy to increase the utility of bioinformatics pipelines.

## Availability and requirements

Project name: REViewer and FlipBook

Project home page: https://github.com/Illumina/REViewer, https://github.com/broadinstitute/flipbook/

Operating systems: REViewer: Linux and macOS; FlipBook: Linux, macOS, and Windows

Programming languages: C++ (REViewer) and Python (FlipBook)

License: GNU GPLv3 (REViewer) and MIT (FlipBook)

## Abbreviations

REViewer: Repeat expansion viewer
STR: Short tandem repeat
In-repeat read: Read fully contained in the repeat sequence
TP-PCR: Triplet primed PCR
EH: ExpansionHunter

## Acknowledgements

Rajan-Babu IS, Law HY, Yoon CS, Lee CG, Chong SS. Simplified strategy for rapid first-line screening of fragile X syndrome: closed-tube triplet-primed PCR and amplicon melt peak analysis. Expert Rev Mol Med. 2015 May 4;17:e7. doi: 10.1017/erm.2015.5, reproduced with permission.

## Supplementary information

Additional file 1

Note S1: Description of the concordance study dataset, Note S2: Description of the wrapper script, Table S2: REViewer allele structure of *FMR1* reference samples

Additional file 2

Table S1: A description of 133 repeats used for the concordance study. Each row describes the basic sample/repeat information (columns Sample, Sex, Locus), repeat size estimated by PCR and EH (columns PCR_short, PCR_long, EH_short, EH_long, EH_short_CI, EH_long_CI), size thresholds (columns Premuation, Pathogenic), verdicts based on PCR & EH size estimates (columns PCR_verdict, EH_verdict), and analyst verdicts (columns Analyst 1, …, Analyst 12).

Additional file 3

Figure S1: REViewer pileups and TP-PCR profiles of additional *FMR1* intermediate, premutation, and full-mutation samples (1) NA20232, (2) NA20230, (3) CD00014, (4) GM06892, (5) GM06852, (6) GM07063.

## Notes

https://github.com/Illumina/REViewer

https://github.com/broadinstitute/flipbook

https://broadinstitute.github.io/StrPileups

### Competing Interest Statement

A subset of the authors are or were employees of Illumina, Inc., a public company that develops and markets systems for genetic analysis.

