## Additional file 1 for "REViewer: Haplotype-resolved visualization of read alignments in and around tandem repeats"

### Supplementary information

#### Note S1: Description of the concordance study dataset

Whole genome sequencing for 868 STRs corresponding to 398 subjects were included in the dataset. A total of 793 out of 868 cases are described in (Ibanez et al. 2020), and an additional 75 PCR-validated *FMR1* and *DMPK* repeats came from 100,000 Genomes Project samples. The samples were de-identified and do not correspond to the same IDs as in (Ibanez et al. 2020). Only the 133 STRs whose size confidence interval overlapped or exceeded an intermediate or full expansion threshold were selected for review (**Table S1**).

#### Note S2: Description of the wrapper script

The wrapper script works by creating read pileups for loci of interest in one or many BAM files. As an input, it requires either a folder or comma separated list of aligned and indexed BAM files as well as a regions (BED) file with coordinates of an STR locus and the repeated motif. The script requires access to the reference genome and REViewer, ExpansionHunter, and SAMtools (Danecek et al. 2021) executables which will be used to genotype specified STR loci and create read pileups. Additionally, a custom genotype (such as one determined by other methods) can be specified in the BED file to override ExpansionHunter's estimated genotype. REViewer will then use and adjust read visualizations for this custom genotype. REViewer will then use and adjust read visualizations for this custom genotype.

While running the script, a BED file will be converted into a temporary variant catalog where each region becomes a record in the catalog and which will be used to create read pileups for input file(s). Visualizations in SVG format will be saved into the specified output folder with the option to create a HTML file that contains all results for the run for easy reviewing.

Table S2: REViewer allele structure of *FMR1* reference samples

| Coriell ID | Coriell allele size (genotype) | EH allele size | REViewer allele structure |
|---|---|---|---|
| NA07175 | 23/30 (NL) | 23/30 | $(CGG)_{13}AGG(CGG)_{9}/(CGG)_{10}AGG(CGG)_{9}AGG(CGG)_{9}$ |
| NA06890 | 30 (NL) | 30 | $(CGG)_{10}AGG(CGG)_{9}AGG(CGG)_{9}$ |
| NA07174 | 30 (NL) | 30 | $(CGG)_{10}AGG(CGG)_{9}AGG(CGG)_{9}$ |
| NA07538 | 29/29 (NL) | 29/29 | $(CGG)_{9}AGG(CGG)_{9}AGG(CGG)_{9}/(CGG)_{9}AGG(CGG)_{9}AGG(CGG)_{9}$ |
| NA07541 | 29/31 (NL) | 29/31 | $(CGG)_{9}AGG(CGG)_{9}AGG(CGG)_{9}/(CGG)_{10}AGG(CGG)_{10}AGG(CGG)_{9}$ |
| NA20243 | 29/41 (NL) | 29/32 | $(CGG)_{9}AGG(CGG)_{9}AGG(CGG)_{9}$/could not be ascertained |
| NA20238 | 29/30 (NL) | 29/30 | $(CGG)_{9}AGG(CGG)_{9}AGG(CGG)_{9}/(CGG)_{9}AGG(CGG)_{9}AGG(CGG)_{10}$ |
| NA20244 | 41 (NL) | 40 | $(CGG)_{9}AGG(CGG)_{8}AGG(CGG)_{21}$ |
| NA20234 | 31/46 (IM) | 31/46 | $(CGG)_{10}AGG(CGG)_{9}AGG(CGG)_{10}/$ $(CGG)_{9}AGG(CGG)_{9}AGG(CGG)_{13}AGG(CGG)_{12}$ |
| NA20232 | 46 (IM) | 45 | $(CGG)_{9}AGG(CGG)_{35}$ |
| NA20235 | 29/45 (IM) | 29/40 | $(CGG)_{9}AGG(CGG)_{9}AGG(CGG)_{9}/(CGG)_{10}AGG(CGG)_{29}$ |
| NA20236 | 31/53 (IM) | 31/44 | $(CGG)_{10}AGG(CGG)_{9}AGG(CGG)_{10}/(CGG)_{44}$ |
| NA20230 | 53 (IM) | 65 | $(CGG)_{65}$ |
| CD00014 | 56 (PM) | 58 | $(CGG)_{9}AGG(CGG)_{9}AGG(CGG)_{38}$ |
| NA20231 | 76 (PM) | 82 | $(CGG)_{10}AGG(CGG)_{71}$ |
| NA20242 | 30/73 (PM) | 30/67 | $(CGG)_{10}AGG(CGG)_{9}AGG(CGG)_{9}/(CGG)_{9}AGG(CGG)_{9}AGG(CGG)_{47}$ |
| NA06892 | 93 (PM) | 73 | $(CGG)_{10}AGG(CGG)_{62}$ |
| NA20240 | 30/80 (PM) | 95/95 | could not be ascertained/could not be ascertained |
| NA06907 | 29/85 (PM) | 29/95 | $(CGG)_{9}AGG(CGG)_{9}AGG(CGG)_{9}$/could not be ascertained |

| | | | |
|---|---|---|---|
| NA06896 | 23/95-120-140 (PM) | 23/80 | $(CGG)_{13}AGG(CGG)_{9}/(CGG)_{10}AGG(CGG)_{69}$ |
| NA06891 | 118 (PM) | 110 | $(CGG)_{110}$ |
| NA07862 | 501-550 (FM) | 99 | $(CGG)_{99}$ |
| NA07294 | n.a (FM) | 102 | $(CGG)_{102}$ |
| NA04025 | 645 (FM) | 112 | $(CGG)_{111}$ |
| NA20239 | 20/183-193 (FM) | 20/103 | $(CGG)_{10}AGG(CGG)_{9}/(CGG)_{103}$ |
| NA07063 | n.a (FM) | 32/93 | $(CGG)_{9}AGG(CGG)_{22}/(CGG)_{93}$ |
| NA06852 | >200 (FM) | 81 | $(CGG)_{10}AGG(CGG)_{70}$ |
| NA06897 | 477 (FM) | 80 | $(CGG)_{10}AGG(CGG)_{69}$ |
| NA07537 | 28-29/>200 (FM) | 29/72 | $(CGG)_{9}AGG(CGG)_{9}AGG(CGG)_{9}$/could not be ascertained |

NL, normal; IM, intermediate; PM, premutation; FM, full-mutation; n.a, not available; EH, ExpansionHunter
Allele size includes AGG interruptions
