## Supplementary figures and images for "REViewer: Haplotype-resolved visualization of read alignments in and around tandem repeats"

### Additional file 3

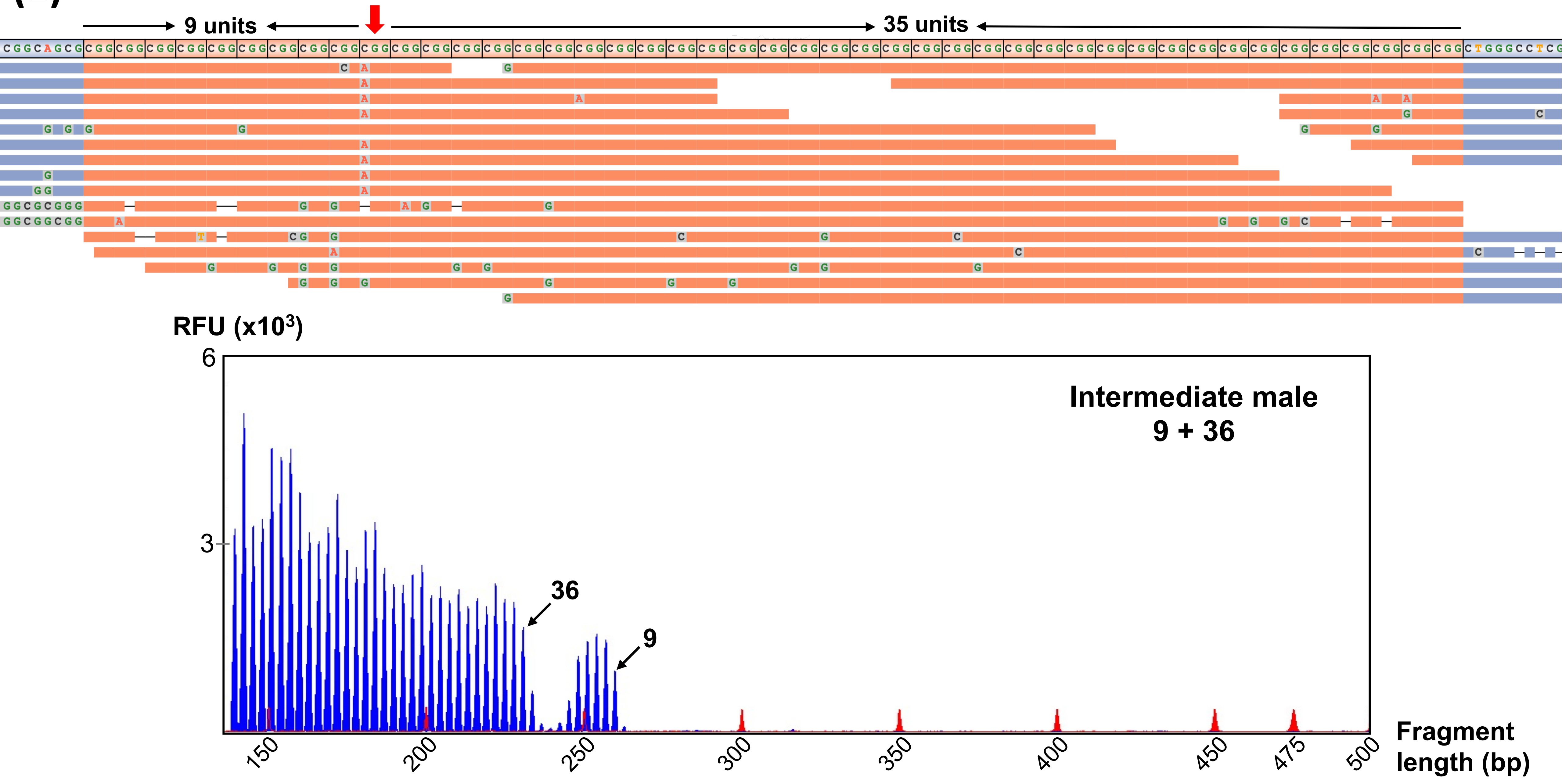

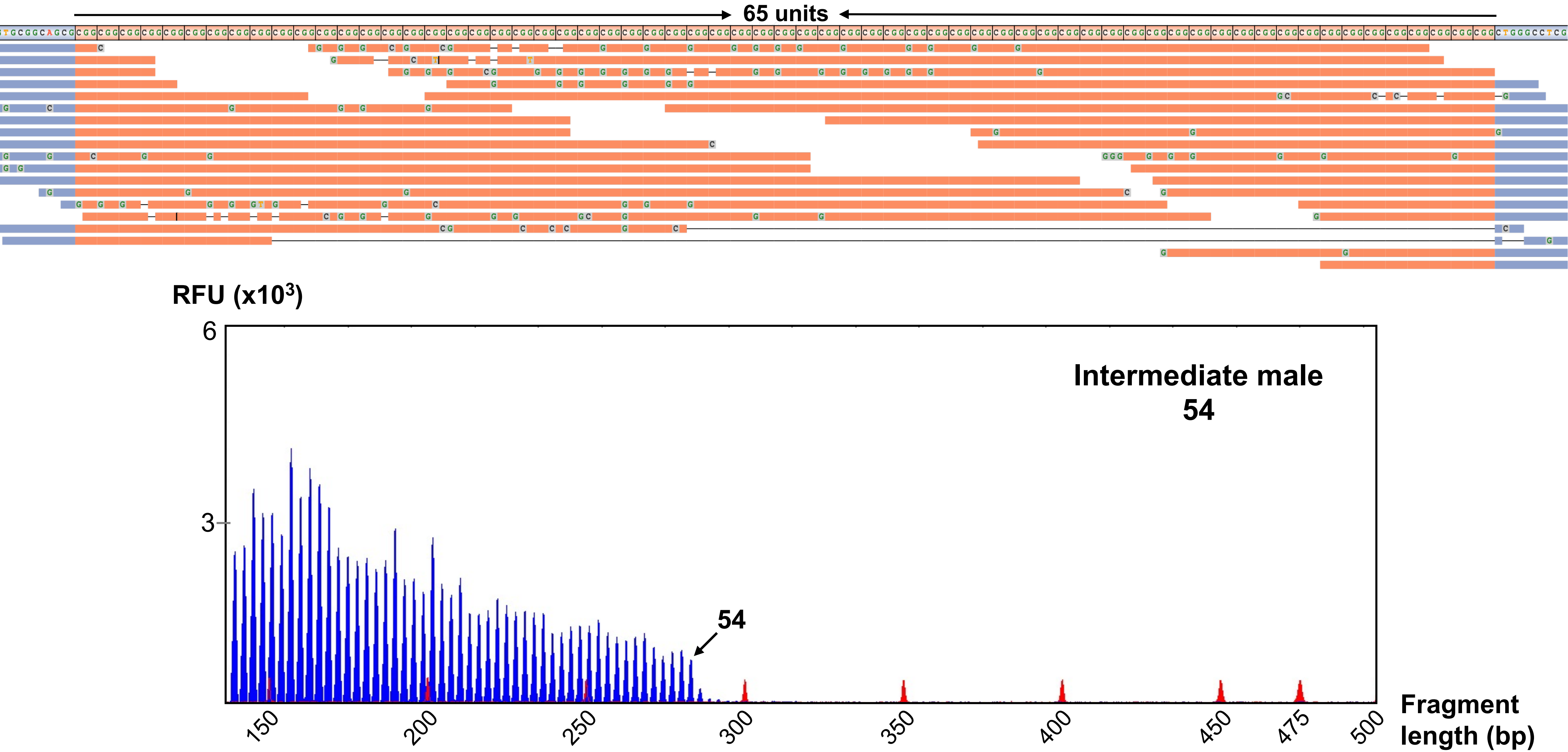

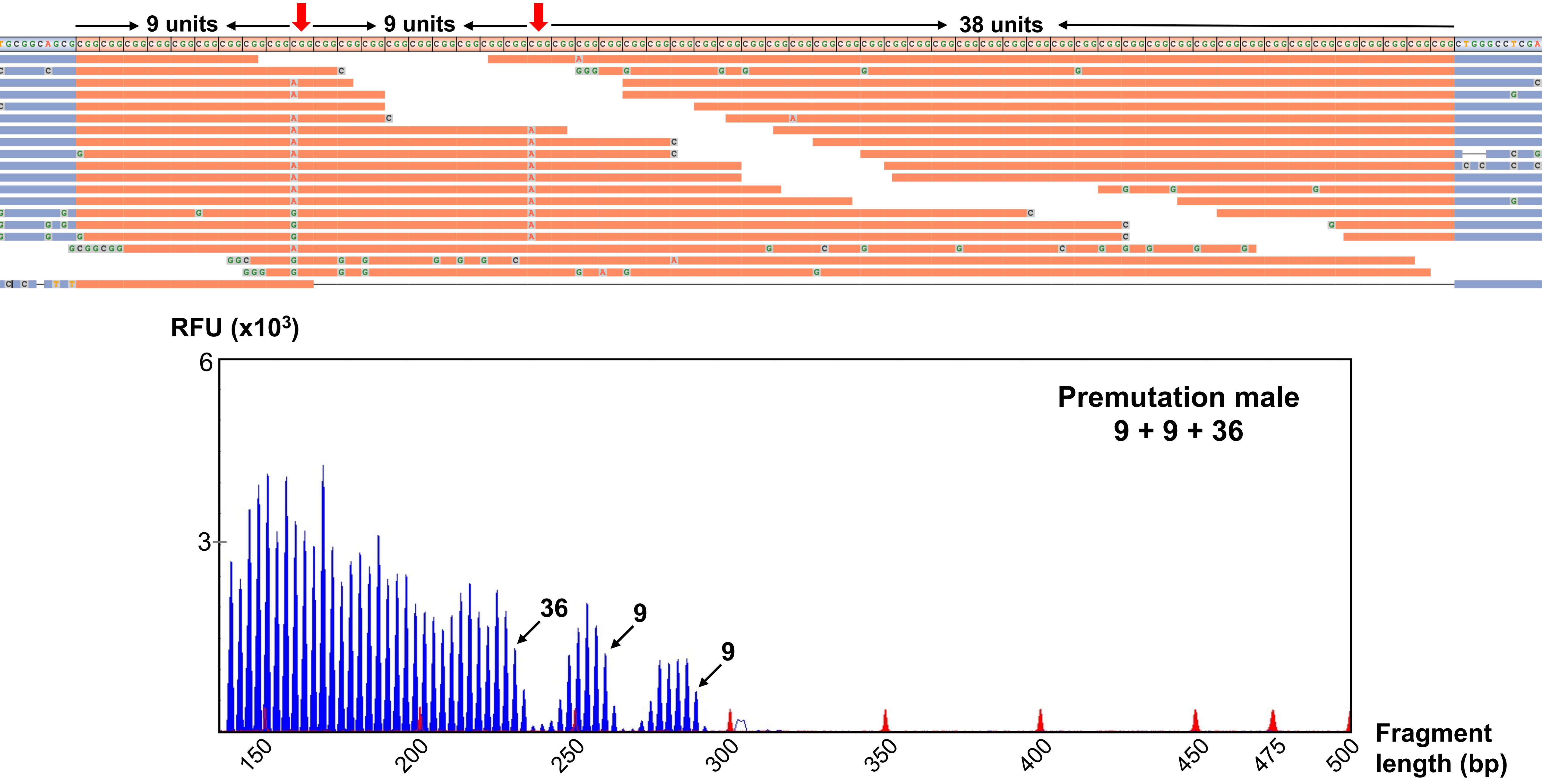

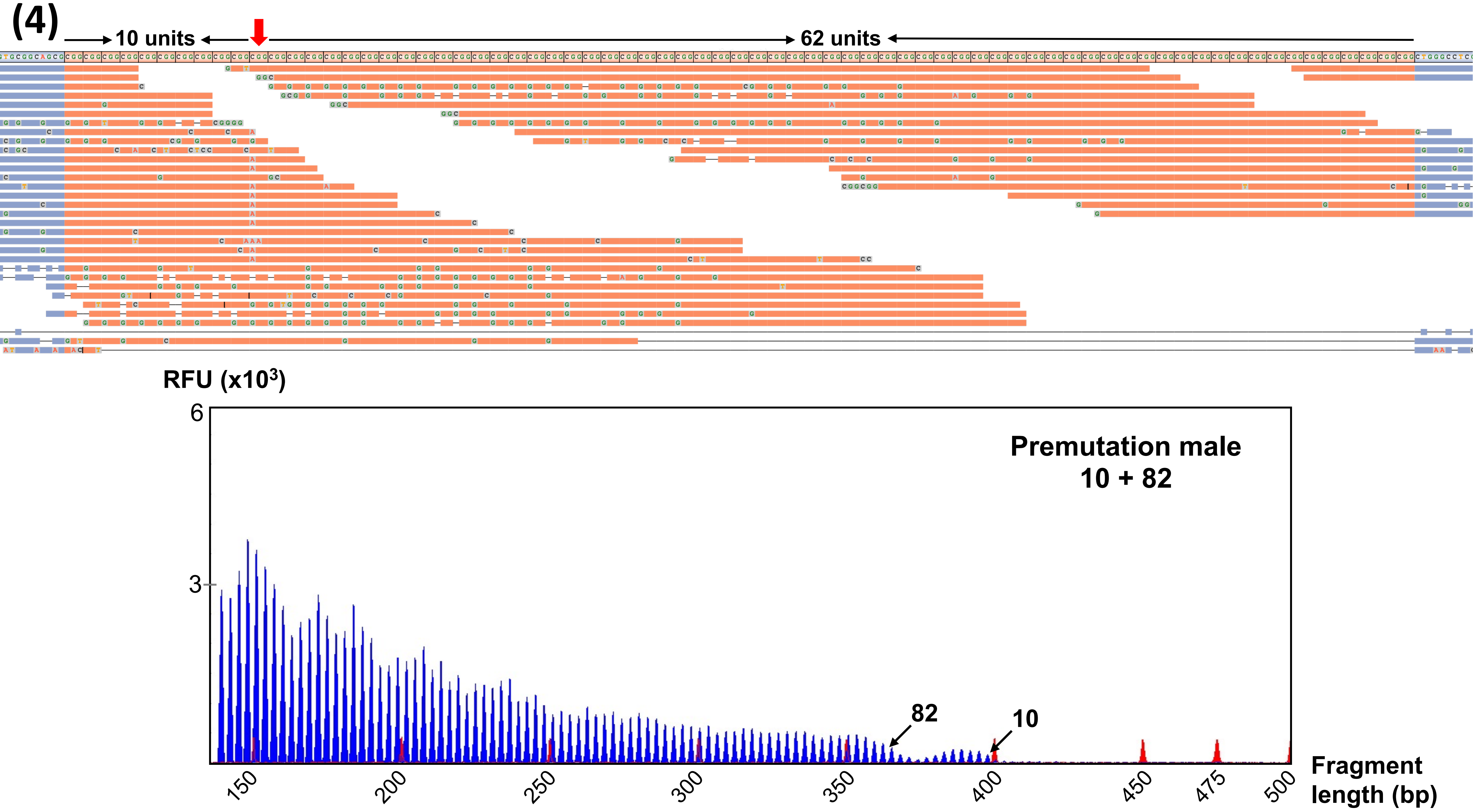

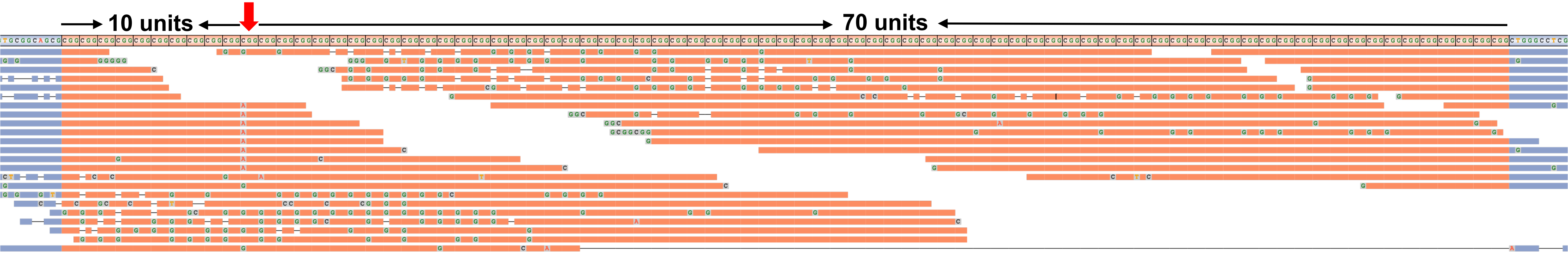

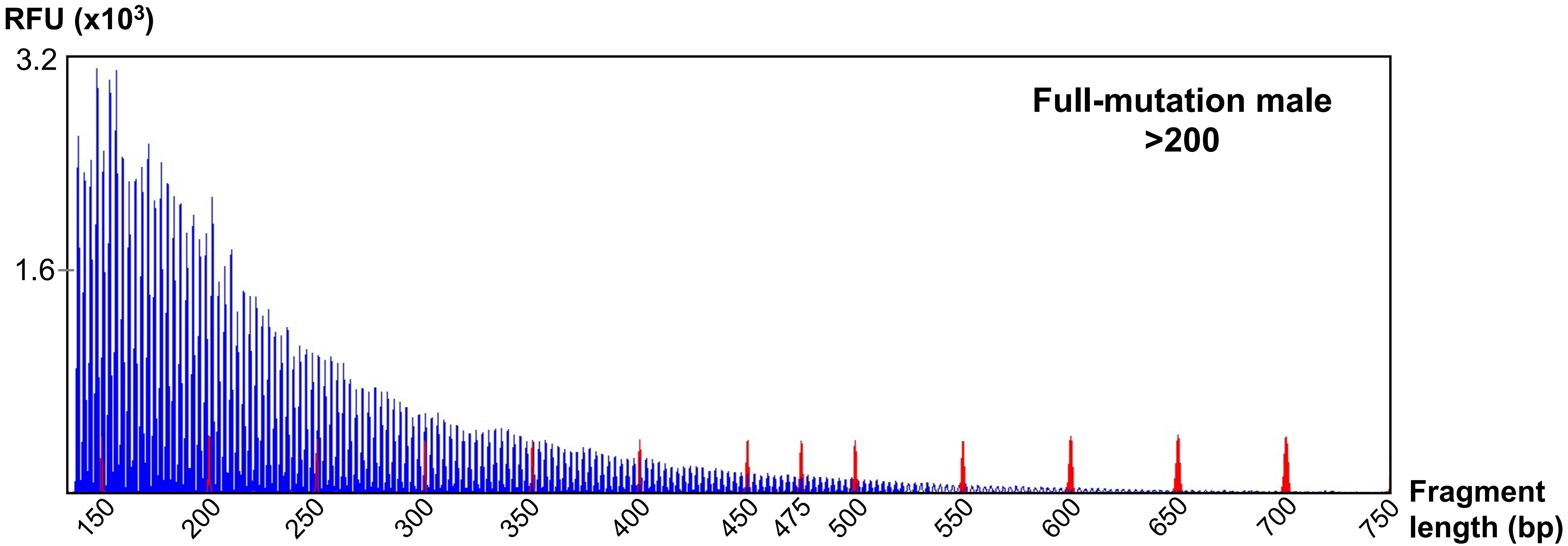

(6a)

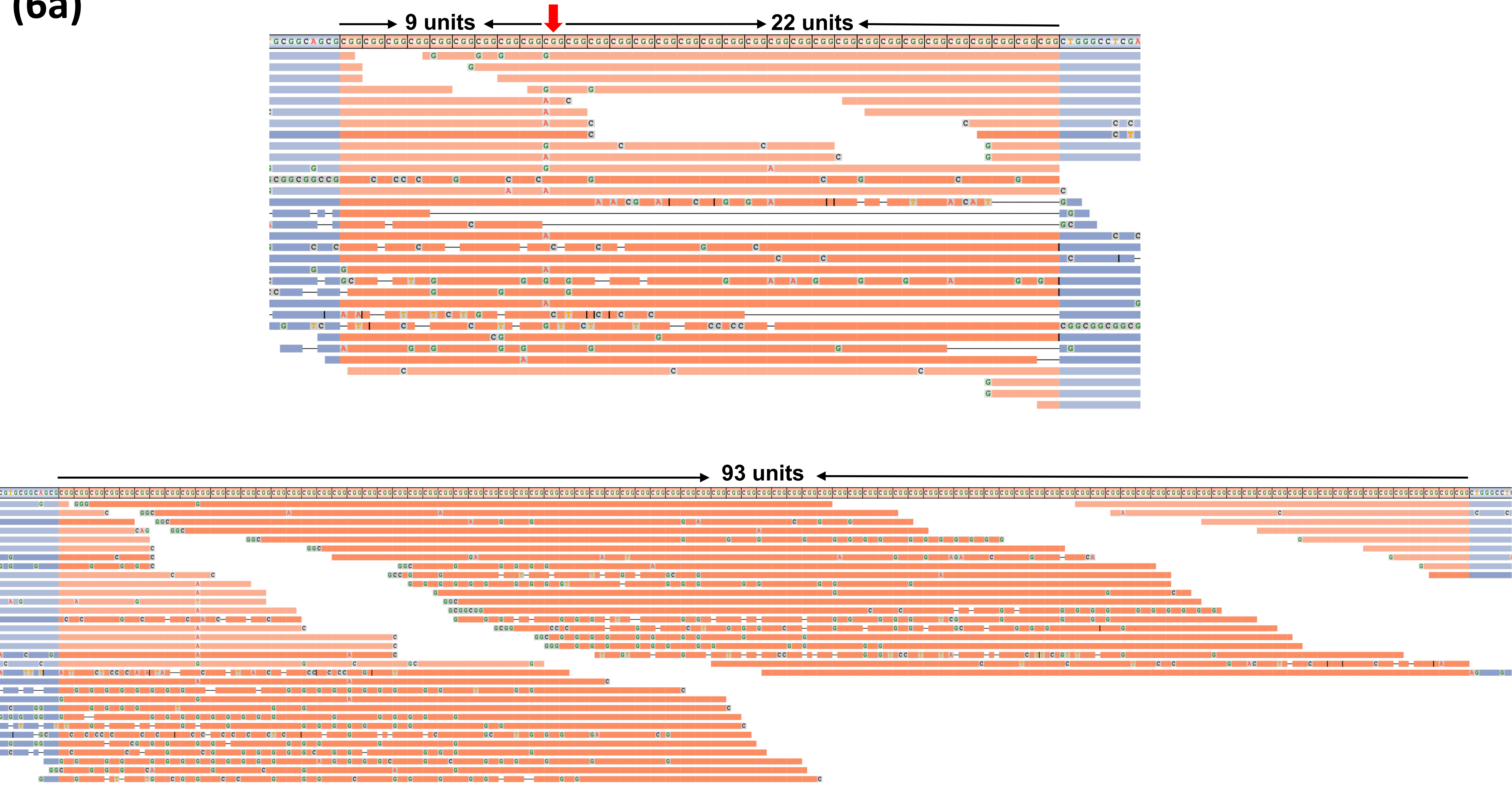

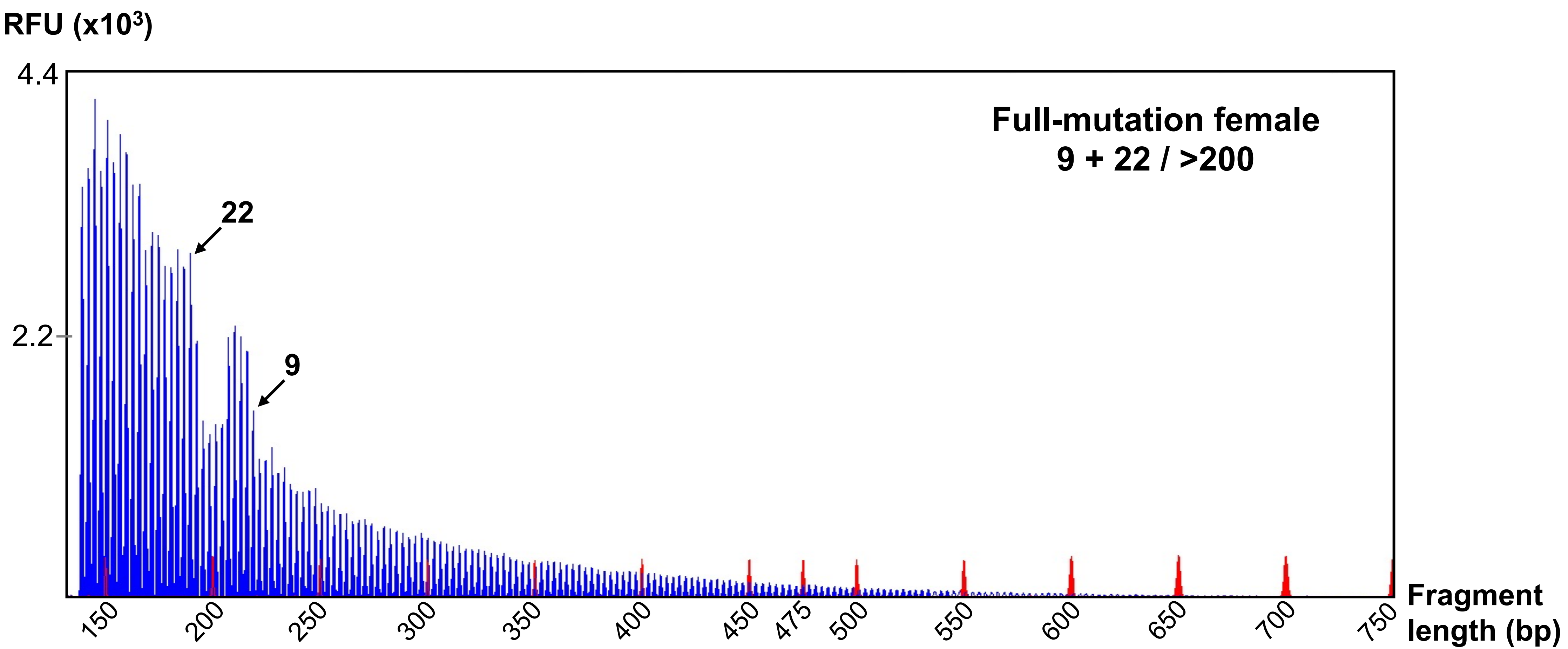
